# Peer presence and familiarity as key factors to reduce cocaine intake: evidence from translational research

**DOI:** 10.1101/286518

**Authors:** E Giorla, S Nordmann, Y Pelloux, P Roux, S Rosellini, K Davranche, C Montanari, Giorgi Lisa, A Vilotitch, P Huguet, P Carrieri, C Baunez

**Author notes:** the authors contributed equally.

## Abstract

Stimulant use, including cocaine, is a major public health issue and decreasing intake can reduce associated harms. We used a translational research approach (experimental for rats and observational for humans) to explore the influence of peer presence and familiarity on the frequency of self-administered cocaine. In both rats and humans, we compared cocaine intake when alone with intake when peers with different characteristics (familiar or not, cocaine-naive or not, dominant or subordinate) were present. In both rats and humans, the risk of drug consumption was reduced when a peer was present and further diminished when the peer was unfamiliar (vs familiar). In rats, the presence of a cocaine-naive peer further decreased cocaine consumption.

The presence of a non-familiar and drug-naive peer represents key conditions to diminish cocaine intake. Our results indirectly support the use of social interventions and harm reduction strategies, in particular supervised consumption rooms for stimulant users.

## INTRODUCTION

Drug use often occurs in a social environment that can influence drug consumption patterns and related behaviors. A social environment encompasses two types of factors: distal social factors (i.e. present in an individual’s broader social environment, but may not be immediately present when drug use occurs) and proximal social factors (i.e., immediately present at the time of drug use).

The influence of distal social factors on drug use has been widely studied. Stress, isolation and rejection are associated with higher rates of drug use in humans ^1,2^ and in animal models ^3–5^. In contrast, strong familial ties are associated with lower rates of drug use in humans ^6–9^,while an enriched environment for animals is associated with lower rates in rodents and monkeys ^3,5,10^. In humans, epidemiological studies have shown that social network characteristics in people who use drugs are major determinants of drug use initiation ^11,12^,persistence ^13,14^, increase ^15^ and cessation ^16,17^. They are also major determinants of risk practices, such as sharing injecting equipment ^18^.

To our knowledge, no study to date has specifically focused on the influence of peer presence, peer characteristics and peer familiarity and their effect on cocaine consumption in humans. The few existing studies in this area only examined the influence of peer presence and close relationships on outcomes such as alcohol use ^19^ and craving during stressful events ^20^.

With respect to animals, studies examining proximal social factors and drug use are relatively recent and suggest the following: 1) social contact and drug use are both rewarding, although social contact may outweigh the rewarding effect of drug use ^21,22^ 2) the presence of peers influences drug consumption ^23^, 3) this influence is substance-specific ^24^ and, 4) whether or not the peer is also self-administering a drug can differentially influence the self-administration behavior ^25^. However, in these animal studies, many characteristics of the peers such as familiarity, dominance status, former experience of the drug, were not investigated.

The influence of proximal social factors is of particular interest when exploring the use of stimulants such as cocaine for several reasons. First, the use of stimulants, in particular cocaine, is of major clinical importance due to the lack of effective pharmacologic treatment. Second, cocaine (and stimulants in general) is characterized by a short half-life, which can lead to very frequent consumption. This in turn amplifies the risk of infection (HIV, hepatitis) and associated complications, and raises wider public health concerns. Therefore, understanding how to decrease stimulant frequency is essential to reduce stimulant-associated harms, and may help stakeholders develop effective social and harm reduction interventions to reduce the associated burden.

To date, research on the influence of proximal social factors in animal and human studies remains sparse. This is perhaps due to the difficulty of “translating” certain social contexts from rats to humans and vice versa, and of analyzing data in a standardized fashion to make comparison possible.

This is the first study to propose a translational approach for both data collection and statistical modeling, in order to explore the extent to which peer presence, peer relationship (familiarity and dominance/subordination) and peer drug exposure history, can influence drug consumption in rats and humans self-administering cocaine.

### Translational issues

The challenges in combining the results of the two studies were overcome thanks to several multidisciplinary sessions which included epidemiologists, neurobiologists/psychologists and statisticians. Working together, these stakeholders ensured the validity of both studies’ designs, and decided “a priori” on the strategic statistical analysis plan to implement in order to facilitate comparison.

More specifically, the three challenges overcome concerned: 1) design 2) explanatory variables 3) outcomes and statistical analysis.

With respect to the first challenge concerning design, it should be noted that cocaine administration can be randomized in rats but not in humans, at least in France. We therefore used a randomized plan (see experimental design section for rats) for the former, but for ethical concerns, an observational cross-sectional epidemiologic design for the latter. This observational design allowed us to retrospectively explore each episode of cocaine/methylphenidate consumption and the associated social context. We adapted a methodology already used in network analysis by Buchanan and Latkin ^16^. It is important to underline that the experimental conditions for humans (i.e., intake in the presence or not of peers, intake with or without familiar, subordinate or cocaine-naÏve peers) mirrored the experimental conditions in the study for rats.

The second major challenge was to “translate” the concepts of familiarity, subordination among peers and cocaine naivety from rats to humans. Due to the lack of literature on this specific issue, this “translation” was constructed using open-ended questions which explored these same three concepts. This approach helped to provide an operational definition of what familiarity, subordination, and cocaine naivety mean for cocaine-using peers.

For the third challenge, we used the same outcome (“frequency of cocaine injection/duration of the session/episode”) and the same models in both studies to provide both an adequate estimate of the association between each “social context” and the outcome. Poisson regression models (based on generalized estimating equations for humans and mixed models for rats) were used to obtain comparable estimates of the associations found between each social context and the frequency of cocaine intake in rats vs. humans. These models provide, for each predictive/explanatory variable associated with the frequency of cocaine/methylphenidate intake, an estimate of the incidence rate ratio (IRR) or relative risk and its 95% confidence intervals. Confidence intervals not containing 1 indicate a significant association. IRRs are a measure of the association between the explanatory variable and the frequency of cocaine intake. For example, a significant IRR of 0.30 for the presence of a peer compared with being alone means that the relative risk of intake of cocaine over the duration of the episode decreases by 70% when an individual (rat or human) is with a peer. The use of these IRRs allows comparison of the association found between each social context and the frequency of cocaine use in both rats and humans.

## RESULTS

A translational and transdisciplinary research approach (experimental in rats – observational in humans) using Poisson regression models was used to explore the influence of peer presence and peer relationship on subjects self-administering cocaine (outcome). In both populations, cocaine intake was compared when they were alone and when they were with peers having different characteristics (familiar or not, cocaine naive or not, dominant or subordinate). Due to the observational nature of the human study and the presence of confounding for the 77 individuals enrolled, the analyses were also adjusted for potential correlates and confounding variables.

### Human study

#### Description of Human Study Group’s Characteristics

The median [interquartile range (IQR)] frequency of stimulant consumption was 1.05 [0.05-2.6] when subjects were alone and 1 [0.3-2] when with peers. Two thirds of the study sample reported either cocaine use alone or with methylphenidate.

Table 1 describes participant characteristics (N=77) and details of drug use episodes involving intranasal and intravenous routes of administration (246 episodes).

**Table 1.** HUMANS-Characteristics of the study sample (N=77) and drug use at any episode reported by participant (N=246) ^1^including alcohol ^2^AUDIT-c score ≥3 for women and ≥4 for men *the denominator may be lower due to missing information for some variables (always below 10%)

|  | Study sample<br>N=77 |
| --- | --- |
| Male gender | 83.1 |
| Age – years | 41 [34; 47] |
| Daily stimulant use | 36.4 |
| Age at first drug use | 17 [15 ; 22] |
| >20 cigarettes per day | 44.7 |
| Hazardous alcohol use<sup>2</sup> | 48.7 |
| Stable housing | 41.6 |
| Financial problems | 66.2 |
| High school certificate | 32.0 |
|  | Study sample<br>N=246* |
| **Peer presence** |  |
| Alone | 26.8 |
| Presence of one peer | 48.8 |
| Presence of a group (2 or more peers) | 24.4 |
| **Familiarity** |  |
| Alone | 26.9 |
| Familiar peer | 29.4 |
| Non-familiar peer | 19.2 |
| Presence of a group (two or more peers) | 24.5 |
| **Route of drug administration** |  |
| Sniffing | 32.9 |
| Injection | 67.1 |
| **Private (vs Public) site when using drugs** | 39.6 |
| **Time of drug use** |  |
| Morning | 26.6 |
| Afternoon | 42.2 |
| Evening/Night | 31.1 |

<sup>1</sup>including alcohol
<sup>2</sup> AUDIT-c score $\geq 3$ for women and $\geq 4$ for men

More than 80% of the study sample were males with median age of 41 years. By definition they were all cocaine users and more than one third reported daily use. Median age at first use was 17 years. Most reported polysubstance use, approximately half were classified as hazardous alcohol users and 45% as heavy smokers (more than 20 cigarettes/day). One third reported having a high school certificate and 42% declared stable housing. The majority (66.2%) reported financial problems.

This study group reported 246 episodes (on average 3.2 episodes per person) of stimulant use alone (26.8%), with a peer (48.8%) or in a group (24.4%). The majority of episodes reported by the study group occurred in presence of a familiar peer (29.4%), followed by alone (26.9%) and within a group (24.5%) while episodes of stimulant use alone accounted for 19.2%. It is worth noting that in 67.1% of the reported episodes, study participants had injected the drug, and that most episodes of stimulant use occurred in a public space. The majority of episodes occurred in the afternoon and the median number of substances used (including alcohol) at any episode was 2.

### Rat and Human studies

#### Influence of peer presence

Rats in the “alone” condition took an average of 13 (+/- 1.8) cocaine injections during the one-hour self-administration session. When a peer was present (familiar or not), the average value was 10 (+/- 1.6) (Fig. 1A). We observed a 51% increase in the relative risk of cocaine intake when rats were alone, compared with when they were in the presence of a peer (IRR[95%CI]=1.51[1.41-1.64], p<0.0002). (Fig. 1B).

**Fig1A.**
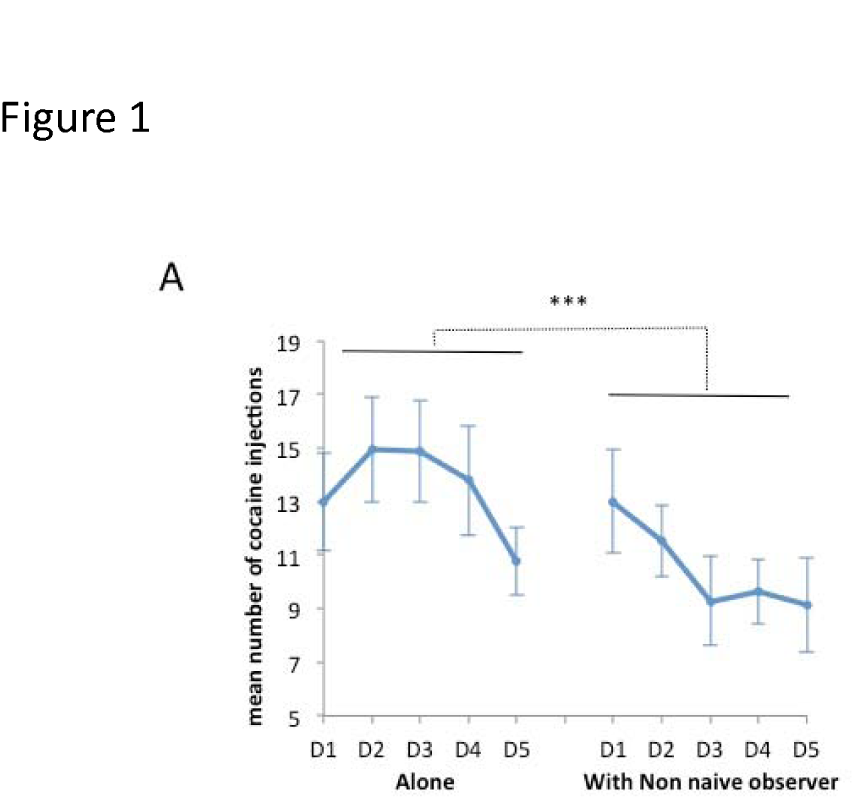
Influence of the presence of a peer on cocaine self-administration in rats. The results are illustrated as the mean (± SEM) number of cocaine injections (80µg/90µl/injection) per 1h-session during 5 consecutive sessions of baseline (“alone”, D1 to D5) and during 5 consecutive sessions in presence of a peer (“with non-naive observer”, D1 to D5, n=14). ***: p<0.0001 GLM analysis

We observed a comparable effect when a present peer also consumed cocaine: when they were alone, rats had a 3 5 % increased risk of consuming cocaine (IRR[95%CI]=1.35[1.25-1.46] compared with when they were with peers also consuming cocaine (see Fig. 1B for levels of consumption).

**Fig1B.**
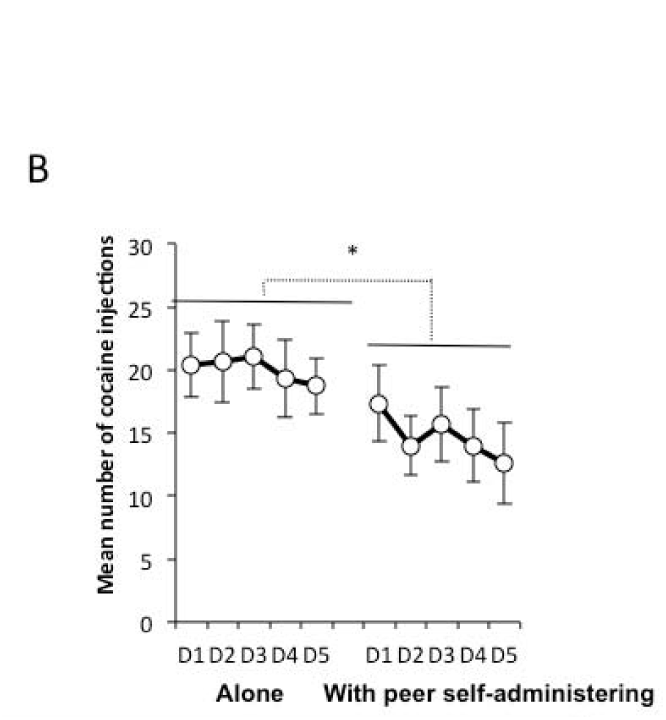
Influence of the presence of a peer self-administering cocaine on cocaine self-administration. The results are illustrated as the mean (± SEM) number of cocaine injections (80µg/90µl/injection) per 1h-session during 5 consecutive sessions of baseline (“alone”, D1 to D5) and during 5 consecutive sessions in co-administration (“with another self-administrating rat”, D1 to D5, n=14). ***: p<0.05 GLM analysis

In the human study, results showed a 59% increased relative risk of cocaine intake during an episode when alone with respect to when one peer was present (IRR[95%CI]=1.59[1.07; 2.38], p=0.023), and an 80% (statistically significant) increased relative risk of cocaine intake when in a group with respect to when only one peer was present (IRR[95%CI] =1.80[1.31; 2.47], p=0.0003) (Table 2 and Fig. 1C).

**Table 2.**
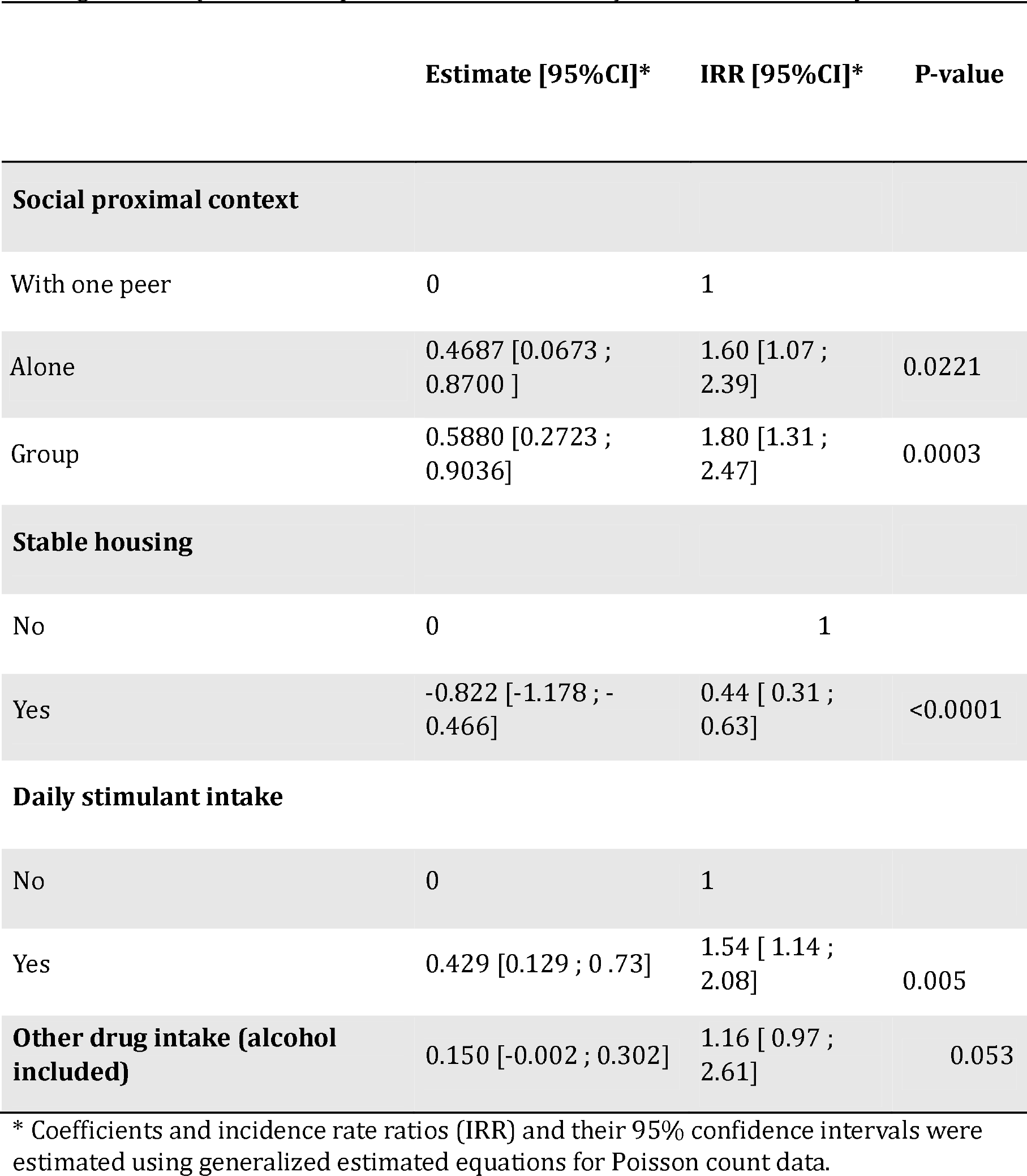
Human study-Association between peer presence and cocaine use frequency among humans (N=77, 246 episodes of stimulant use) – multivariable analysis. * Coefficients and incidence rate ratios (IRR) and their 95% confidence intervals were estimated using generalized estimated equations for Poisson count data.

|  | Estimate [95%CI]* | IRR [95%CI]* | P-value |
| --- | --- | --- | --- |
| **Social proximal context** |  |  |  |
| With one peer | 0 | 1 |  |
| Alone | 0.4687 [0.0673 ; 0.8700 ] | 1.60 [1.07 ; 2.39] | 0.0221 |
| Group | 0.5880 [0.2723 ; 0.9036] | 1.80 [1.31 ; 2.47] | 0.0003 |
| **Stable housing** |  |  |  |
| No | 0 | 1 |  |
| Yes | -0.822 [-1.178 ; -0.466] | 0.44 [ 0.31 ; 0.63] | <0.0001 |
| **Daily stimulant intake** |  |  |  |
| No | 0 | 1 |  |
| Yes | 0.429 [0.129 ; 0.73] | 1.54 [ 1.14 ; 2.08] | 0.005 |
| **Other drug intake (alcohol included)** | 0.150 [-0.002 ; 0.302] | 1.16 [ 0.97 ; 2.61] | 0.053 |

**Fig1C.**
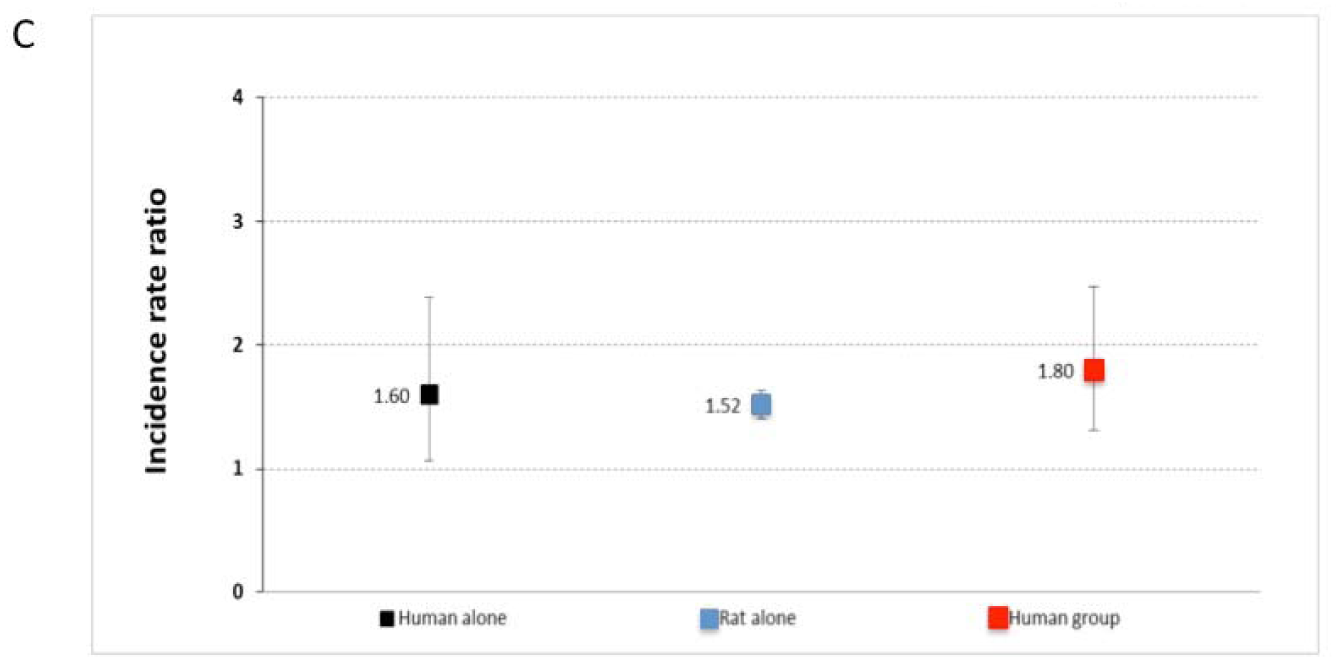
Adjusted incidence rate ratios (relative risk) of frequency of drug consumption depending on the peer presence in humans and in rats. Reference = “presence of one peer”. Black and red squares represent the adjusted incidence rate ratio from the multivariable analysis using GEE Poisson model in humans of the variable peer presence (reference=human with a peer): human alone (black square) and human with group (red square). The lower and upper dashes represent, respectively, the lower and upper bounds of the 95% confidence interval. Blue square represents the adjusted incidence rate ratio from the multivariable analysis using GEE Poisson model in rats of the variable peer presence (reference=rat with a peer). The lower and upper dashes represent, respectively, the lower and upper bounds of the 95% confidence interval.

However, there was no significant difference in the frequency of cocaine intake when a group of peers was present compared with being alone as confidence intervals of IRR overlapped (Table 2).

#### Influence of peer familiarity

In the rat study, the average number of cocaine injections self-administered during the one-hour sessions was lower when the peer was “non-familiar” (7.6 (±0.34)) than when “familiar” (12.7 (±0.24)) (Fig. 2A). The Poisson model analysis showed an increasing relative risk of consumption from the presence of a non-familiar peer (IRR=1, reference group), to that of a familiar peer (IRR[95%CI]: 1.20[1.02-1.42]), and to the condition of being “alone” (IRR[95%CI]: 1.73[1.50-1.98]) (Fig. 2C). It is worth noting that confidence intervals of IRR for “familiar peer” and being “alone” did not overlap. This means that there was a significant difference in drug consumption between “familiar peer” and being “alone”.

**Fig2A.**
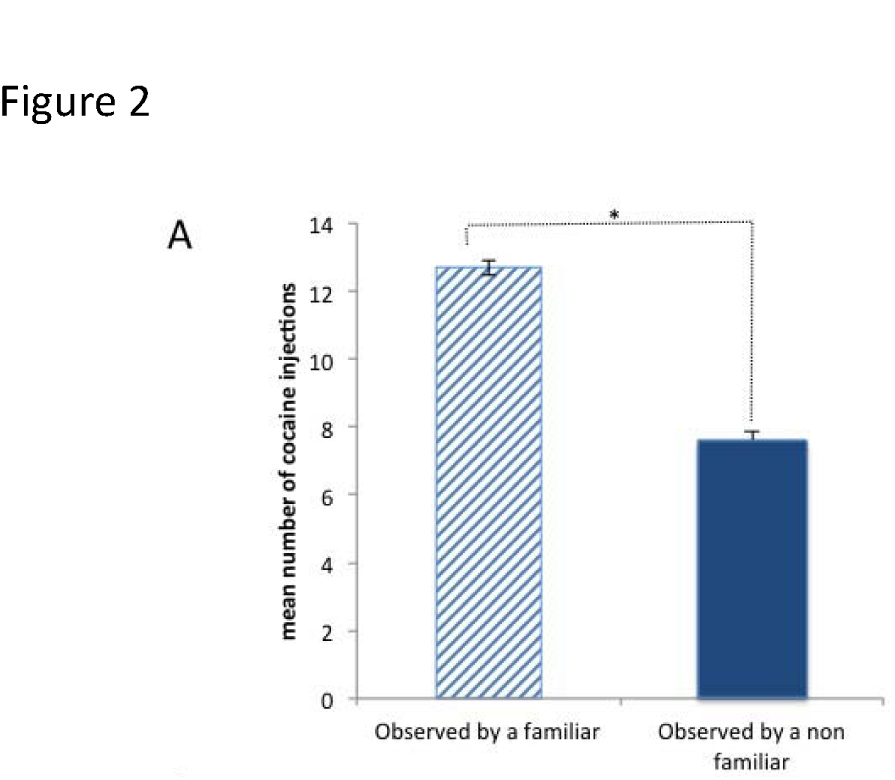
Influence of familiarity with an abstaining peer on cocaine self-administration in rats. The results are illustrated as the mean (± SEM) number of cocaine injections (80µg/90µl/injection) for an average of 5 consecutive days of social interaction with either a familiar peer (striped blue bar, n=8) or an **unknown** peer (“non-familiar”, dark blue bar, n=6).*: p<0.01 GLM analysis.

These results were also confirmed in the second experiment, when rats consumed cocaine in the presence of a peer which also consumed cocaine (see Fig. 2B for number of injections in co-administration within familiar versus strangers): in the mixed Poisson model analysis, we found that compared with being with a non-familiar peer, an increasing relative risk of consumption was observed from being with a “familiar peer” (IRR[95%CI]= 1.10[0.93-1.29]) to being “alone” (IRR[95%CI]=1.41[1.26-1.57]), However, these increases were smaller than those observed in the first experiment.

**Fig2B.**
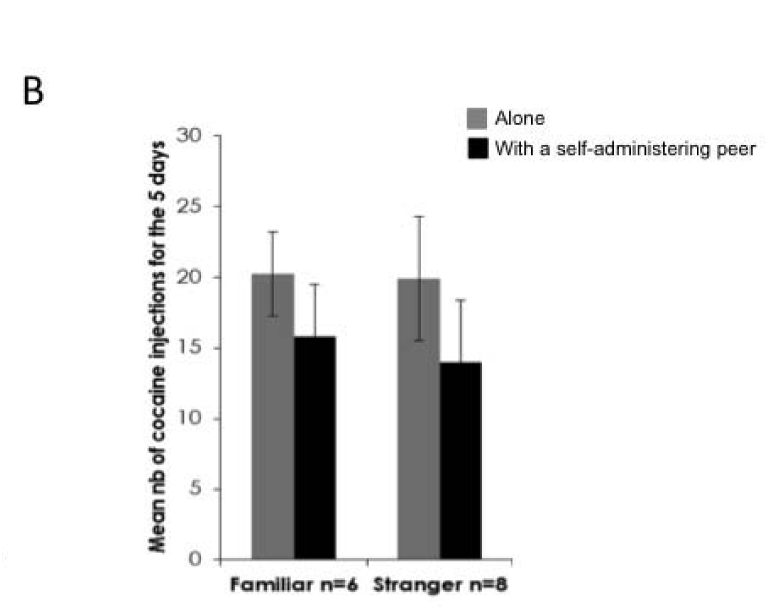
Influence of familiarity with a self-administering peer on cocaine self-administration in rats. The results are illustrated as the mean (± SEM) number of cocaine injections 80µg/90µl/injection) for an average of 5 consecutive days of self-administration when the rat was alone (gray bars) and in presence of a peer also self-administering cocaine (black bars) when this self-administering partner was familiar (left bars, n=6) or a stranger (“non-familiar”, right bars, n=8).

These results are in line with those from multivariable analyses in humans, where an increase in relative risk in stimulant use was observed from non-familiar peer presence (reference category, IRR=1) to familiar peer presence (IRR[95%]=1.62[1.07;2.44]; p-value=0.02), to being alone (IRR[95%]=2.10[1.30;3.41]; p-value=0.003) (Fig. 2C) and to being with a group of peers (IRR[95%]=2.40[1.59;3.61]; p-value<0.0001) (Table 3).

**Table 3.** Human study - Association between familiarity / peer presence and cocaine use frequency among humans (N=77, 246 episodes of stimulant use) – multivariable analysis. * Coefficients and incidence rate ratios (IRR) and their 95% confidence intervals were estimated using generalized estimated equations for Poisson count data.

|  | **Coefficient**<br>**Estimate**<br>**[95% CI]\*** | **IRR [95% CI] \*** | **P-value** |
| --- | --- | --- | --- |
| **Familiarity** |  |  |  |
| Non-familiar peer | 0 | 1 |  |
| Familiar peer | 0.4797 [0.0686 ; 0.8909] | 1.62 [1.07 ; 2.44] | 0.023 |
| Alone | 0.7437 [0.2597 ; 1.2277] | 2.10 [ 1.30-3.41] | 0.003 |
| Group | 0.8741 [0.4654 ; 1.2827] | 2.40 [1.59 ; 3.61] | <.0001 |
| **Unstable housing** |  |  |  |
| No | 0 | 1 | <0.0001 |
| Yes | 0.868 [-1.222 ; -0.513] | 2.38 [1.67 – 3.40] |  |
| **Daily cocaine intake** |  |  |  |
| No | 0 | 1 | 0.002 |
| Yes | 0.468 [0.171 ; 0.766] | 1.60 [1.18 ; 2.15] |  |
| **Number of other substances**<br>**(alcohol included)** | 0.144[-0.007 ; 0.297] | 1.16 [0.99; 1.34] | 0.063 |

**Fig2C.**
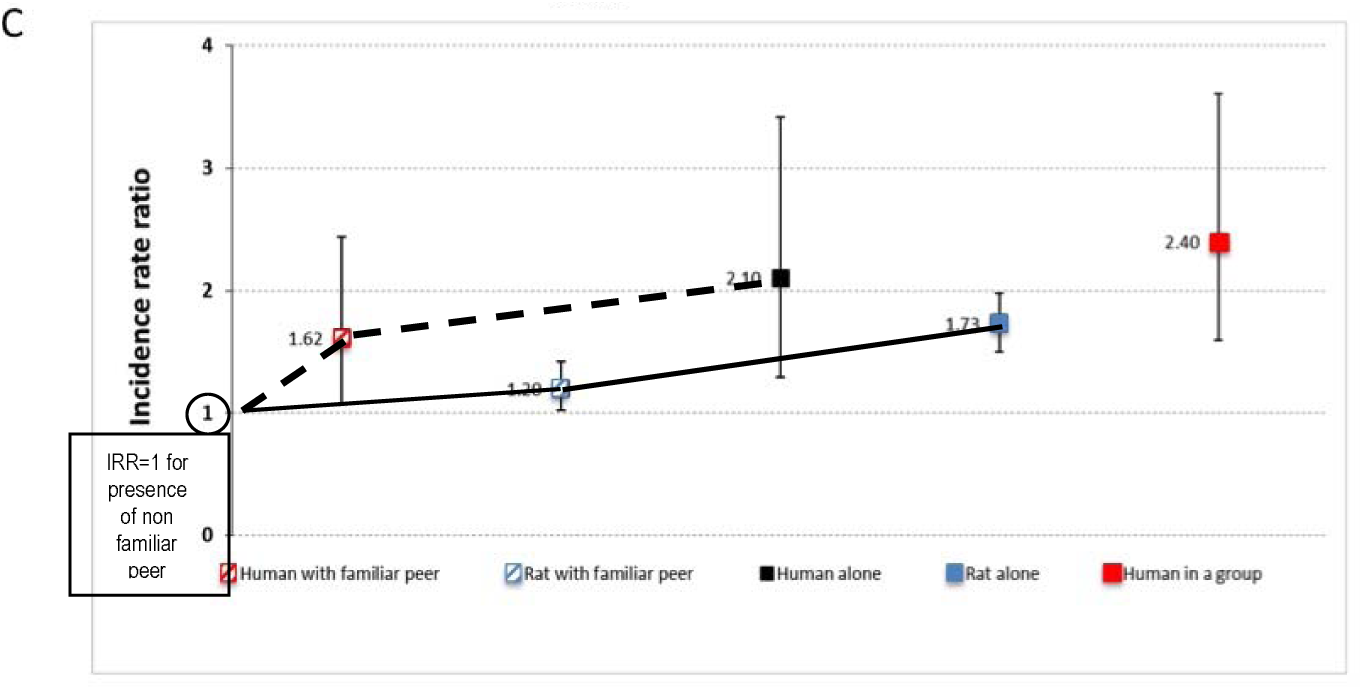
Adjusted incidence rate ratios (i.e. relative risk) for drug consumption depending on peer familiarity in humans and rats. Reference = “non-familiar peer”. Black and red squares represent the adjusted incidence rate ratio from the multivariable analysis using GEE Poisson model in humans of the variable familiarity (reference=human with a non-familiar peer; black square: human alone, red striped square: human with familiar peer, red square: human in group). The lower and upper dashes represent, respectively, the lower and upper bounds of the confidence interval. Blue squares represent the adjusted incidence rate ratio from the multivariable analysis using GEE Poisson model in rats of the variable familiarity (reference=rat with a non-familiar peer; blue striped square: rat with familiar peer, blue square: rat alone). The lower and upper dashes represent, respectively, the lower and upper bounds of the confidence interval.

Other correlates associated with greater frequency of stimulant use in the multivariable analysis were unstable housing (IRR[95%CI]=2.38[1.67; 3.40], p<0.0001), daily stimulant use (IRR[95%CI]=1.60[1.18; 2.15], p=0.002) and number of other substances concomitantly used during the episode (IRR[95%CI]=1.16 [0.99; 1.34], p=0.063).

#### Influence of peer history of drug exposure

Chronologically, in the rat study, after exposure to a non-naive peer, rats were tested again on their own (“alone again”). Cocaine consumption then returned to the baseline level (t=-1.324, p>0.18, Fig.3A).

**Fig3A.**
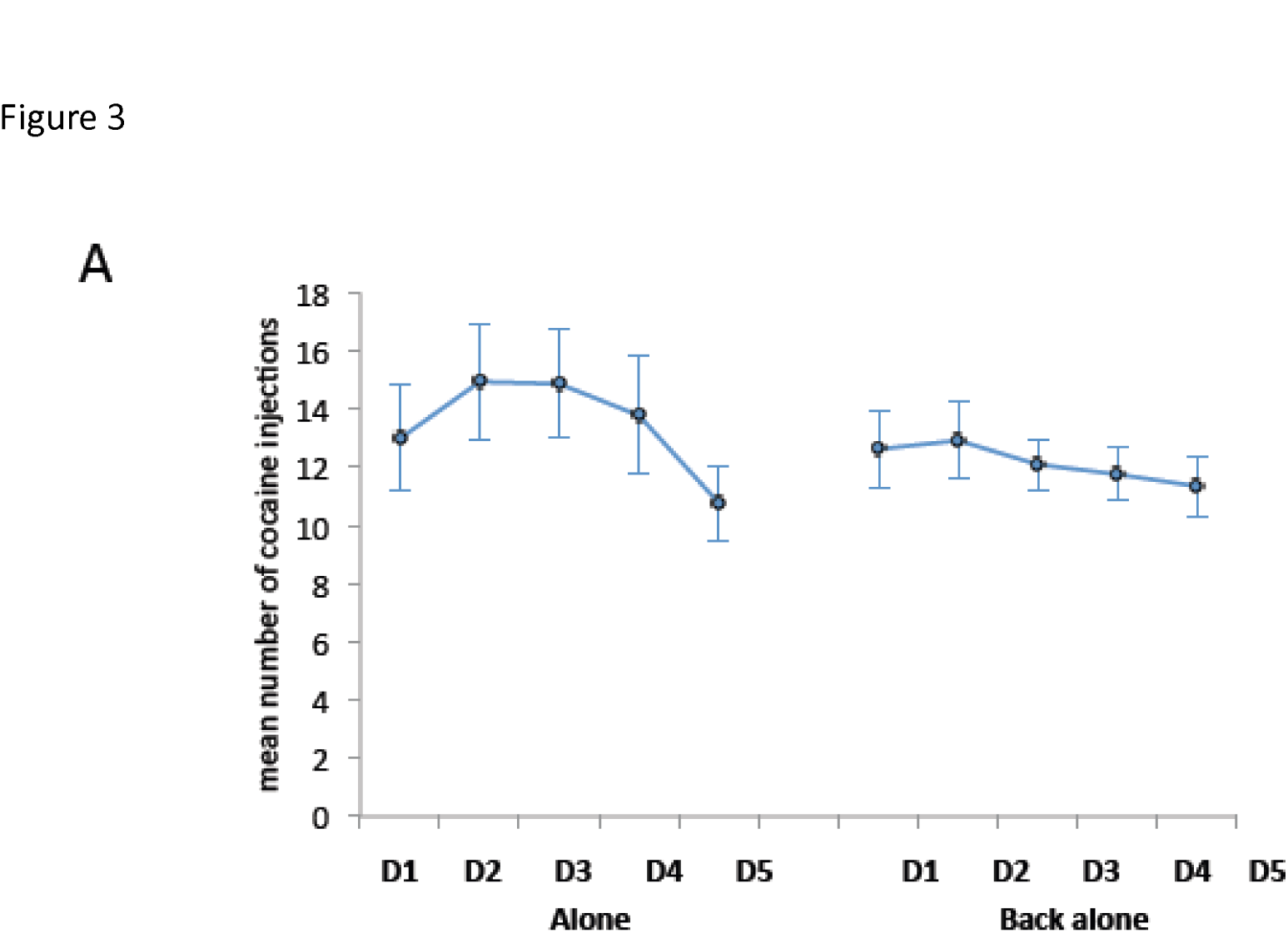
Return to baseline cocaine consumption when rats are tested alone again. The results are illustrated as the mean (± SEM) number of cocaine injections (80µg/90µl/injection) per 1h-session for 5 consecutive days at baseline (“Alone”, D1 to D5: day 1 to 5) and for 5 consecutive sessions alone after the experience of being in the presence of a non-naive peer (“Back alone”, D1 to D5: day 1 to 5).

In the presence of a cocaine-naive peer, rats took an average of 5.6 (±1.2) cocaine injections, and an average of 10.5 (±1.6) injections in the presence of a non-naive peer (familiar and non-familiar included) (Fig. 3B). The Poisson model showed an increasing gradient in the frequency of cocaine consumption from the presence of a naive peer (IRR=1) to the presence of a non-naive peer (IRR[95%CI]: 1.80 [1.57; 2.06], p<0.0001, to the “back alone” (i.e. post-peer) condition (IRR[95%CI]: 2.10[1.84-2.40], p<0.0001), to the “alone” condition (2.31[2.03-2.63], p<0.0001).

In humans, among episodes involving one other peer, we noted that the latter was always a drug user (for episodes with a group, peers were not characterized).

**Fig3B.**
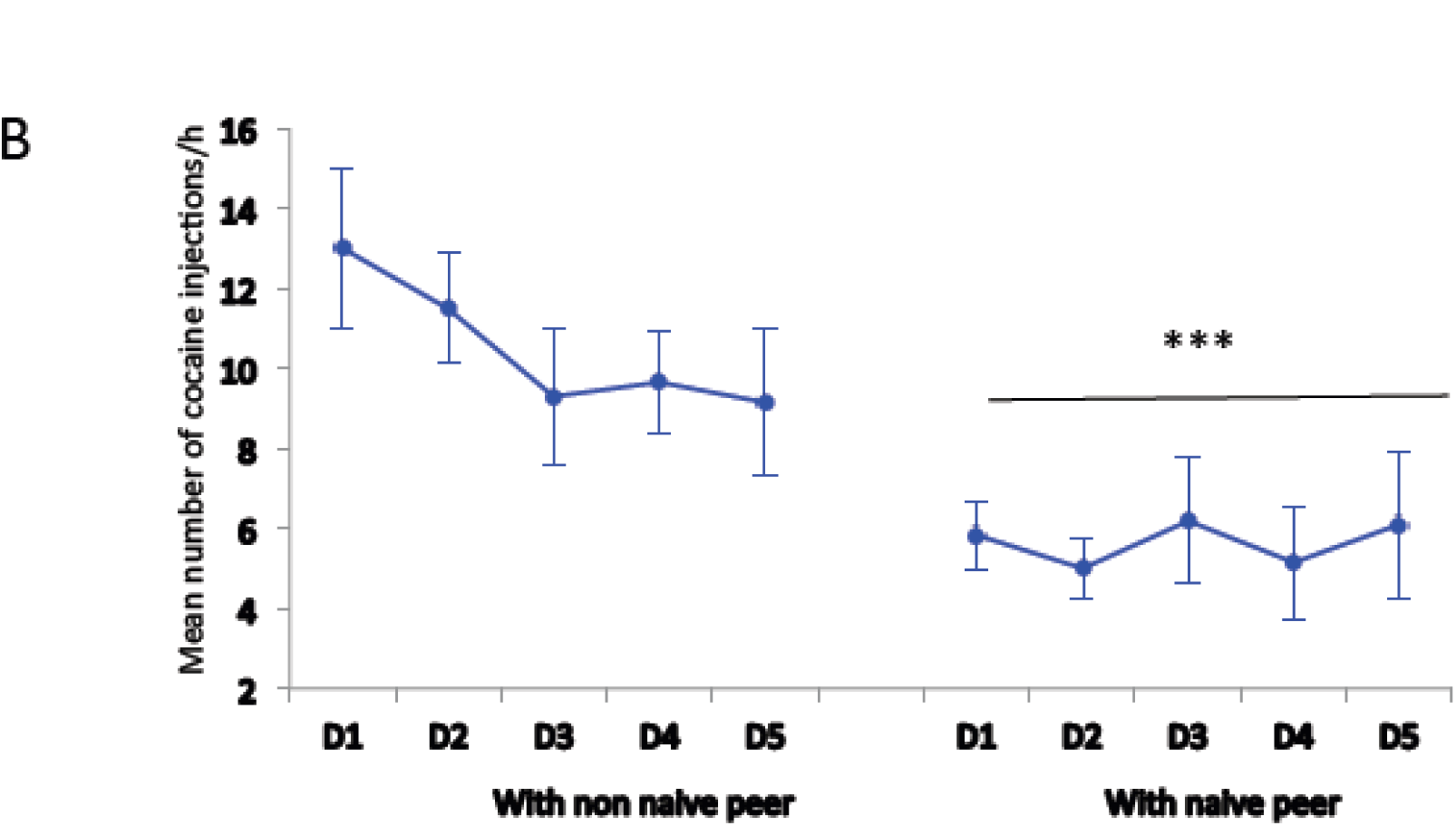
Influence of drug history of the observing peer on cocaine self-administration in rats. The results are illustrated as the mean (± SEM) number of cocaine injections (80µg/90µl/injection) per 1h-session for 5 consecutive sessions under observation of a non-familiar peer with a history of cocaine self-administration (left, “non-naive peer”, D1 to 5: day 1 to 5) and under observation of a non-familiar peer **naive** to cocaine (right, “**naive** peer”, D1 to D5: day 1 to 5) ***: p<0.001 GLM analysis.

#### Dominance/subordination relationship

No significant difference was found when comparing the frequency of cocaine consumption in rats in the presence of a subordinate and in the presence of a dominant familiar peer. Similarly, no significant subordination or dominance effect was found in the analysis of data on humans (in terms of economic dependence and the leader role in drug use contexts). It thus seems that dominance plays a small role in social factors modulating drug use.

## DISCUSSION

This study provides two major contributions. The first is that the influence of proximal social factors on stimulant use is comparable in rats and humans in terms of significance and relative risk. More specifically, our results show that 1) the presence of a peer at the time of stimulant intake has a beneficial effect, in that the frequency of stimulant consumption is reduced; 2) the presence of a non-familiar peer is associated with an even greater decrease in drug use in both human and rats. In other words, in decreasing order, the highest risks of consumption are observed when alone, then with a familiar peer and then with a non-familiar peer; 3) the frequency of cocaine consumption was lower in rats whose peers were cocaine naive than in rats with non-naive peers or rats which were alone (this condition could not be tested in humans). These results were confirmed when rats were in the presence of a peer which also consumed cocaine; 4) no significant effect of peer dominance/subordination was observed in terms of drug consumption.

The consistency of the results found in both rats and humans reinforce their evidence. The second major contribution is that we developed and tested a novel approach in terms of design and statistical analysis to conduct translational research on the influence of proximal social factors on a standardized outcome. This could have important repercussions in research on human behaviors and may encourage other behavioral researchers to adopt a similar approach especially when the research question can be translated into public health actions.

### General influence of social presence

As mentioned above, the main result of this study is that in both humans and rats, cocaine (or stimulant) consumption decreased in the presence of a peer. This confirms previous results in rat-based studies where, just as was the case in our experimental conditions, only one of the two rats used cocaine ^23^. This result therefore supports the hypothesis that the rewarding and reinforcing properties of drugs also depend on whether other individuals are present at the time of drug exposure. Moreover, Thiel and colleagues ^26^ showed in adolescent male rats, that conditioned place preference for nicotine was modified in the presence of a peer. They concluded that the presence of a peer modulates the affective valence of drug use ^26^.

The reduced frequency of cocaine consumption observed here when peers are present can be explained by Fritz and Douglas’ hypothesis that the rewarding effect of social contact ^27^ may outweigh the rewarding effects of drug use ^21,22^. This explanation is concordant with the results of the conditioned place preference test we performed in our rat-based experiment (see supplementary material) where rats (housed in pairs) preferred an environment where they were in contact with their home-cage peer over an environment where they remained alone. This rewarding effect depends however on the dominant/subordinate relationship, as the presence of a dominant peer was not found to be rewarding for the subordinate peer. Since there was no peer dominance/subordination relationship effect on the frequency of stimulant intake in the present study, this rewarding component seems to play a limited role. That said, we were not able to measure this relationship in humans since there was no significant association between being in the company of peer(s) and having a particular mental state (positive, neutral or negative) (data not shown).

Furthermore, in line with our results, an econometric analysis has also shown that reinforcing properties of cocaine diminish when a rat is in presence of an abstaining peer ^28^. This effect is explained by social-learning theories of substance use, which suggest that members of peer groups of PWUD influence each other by imitation ^28^, which could thus be the case in our rat study, but would not explain the human results, since peers were all PWUD as well. The results observed in the presence of a peer also consuming cocaine do not support this hypothesis either. It is interesting to note that this result is the opposite to that previously reported showing increased cocaine intake when both rats were acquiring self-administration simultaneously^29^. The discrepancy might be due to the fact that, in our experiments, the acquisition of drug self-administration was done when rats were alone in the self-administration chamber and the presence of the peer self-administering was only tested after acquisition.

Another explanation could be related to whether or not the peer is under the effect of cocaine at the time of the interaction. It has been shown that adolescent rats increase their ethanol consumption in the presence of an ethanol-intoxicated sibling, but not with a sibling exposed to water or coffee. This suggests that the intoxication status of the peer is an important factor in the way social interaction will affect behavior ^30^. In the first experiment in the present study at the time of the interaction, only the tested rat had access to the drug while the peer had a minimum of 4 hours of abstinence from cocaine, and was therefore regarded as no longer being under the influence of its previous cocaine injections.

### Is social presence a stressor?

Given the well-known association between stress and increased drug intake in both rats and humans ^31^, one might hypothesize that the presence of a peer would increase frequency of cocaine consumption, assuming that an observer could induce some stress. This hypothesis can be ruled out however, since it has been previously shown that in rats and monkeys, the presence of a peer during behavioral execution of a task does not affect blood cortisol levels ^32,33^. Furthermore, the emission of ultrasonic vocalizations (USV) can also be used as a stress indicator. Indeed, stress has been shown to reduce tickling-induced positive 50kHz USV in rats ^34^. In contrast, in the present study, the presence of a peer increased levels of 50 kHz USV (data not shown), while decreasing drug consumption. This suggests that the presence of a peer is not stressful, but rather reinforcing. Since it has been shown that cocaine by itself can induce the emission of positive USV ^35^, our results confirm that increased USV emissions here are more related to the presence of a peer than to cocaine itself.

### Influence of peer familiarity

Our findings show that, in both rats and humans, cocaine consumption was lower when a non-familiar peer was present than when a familiar peer was present. This shows that familiarity is an additional factor that modulates drug consumption. With respect to other types of behavior, in birds, Guillette et al. showed that familiarity with a peer present during a task performance can affect a bird’s behavior ^36^. More specifically, the study showed that in male zebra finches, nest-building skill is socially transmitted only if the demonstrator is a familiar peer and not a stranger. Put briefly, birds use social information, copying the choice only if made by a familiar peer.

### Is a non-familiar peer a stressor?

To return to our study, as discussed above, a stressful situation is usually associated with fewer positive USV in rats. We found this to be true when the peer was not familiar (data not shown). Although non-familiar peers may be considered as creating more stress than familiar peers, the reduced risk of cocaine consumption in our experiment does not argue in favor of a stress effect.

Moreover, in a study examining episodes of stress among humans with substance-use disorders, Preston et al. found that stress events were more likely to occur in situations of social company (interactions with acquaintances, friends, or on the phone) than with family (spouse, child), in places with greater overall activity (bars, outside, walking) and in situations where unexpected experiences occur (interactions with strangers) ^20^.

These results may seem counterintuitive. Indeed, it is known that an emotionally positive context reduces drug consumption ^37^. In our study, rats in the presence of their cage-mate emitted a higher number of positive USV, suggesting an emotionally positive context, although they consumed more cocaine than when in presence of a non-familiar peer. It is known that the primary effect of cocaine is the inhibition of dopamine reuptake, leading to increased extracellular dopamine levels ^38^. Furthermore, it has been shown that the presence of a peer increases extracellular dopamine levels ^33^. If the mere presence of a peer induces a dopamine increase, it might well be possible that a ceiling effect could prevent cocaine from increasing the level further, and this could result in decreased drug efficacy and therefore decreased drug use. The fact that the frequency of cocaine consumption in our study was even lower in the presence of a non-familiar peer would then suggest that extracellular levels of DA can be modulated by levels of familiarity. This remains to be investigated.

Finally, another possible explanation for the difference between the familiar versus non-familiar peer’s influence on self-administration behavior is that the presence of the latter represents a powerful distractor ^39^. A subject’s attention may be focused on the non-familiar peer rather than on the drug, consequently leading to decreased frequency of drug consumption.

In line with this hypothesis, a recent study in monkeys has shown that the presence of a peer increases the activity of attention-related cerebral structures^32^.

### Influence of peer history of drug exposure

The other important result of the present study is that a history of cocaine use in an observer rat induced a higher frequency of drug consumption than the presence of a non-familiar cocaine-naive peer. This result was not likely to be observed in humans, since the peer present during stimulant use was always a drug user.

Although epidemiological studies have highlighted the importance of the relationship within the network between peers at the moment of drug use and on the sharing of injecting equipment ^40^, drug seeking ^19,41–43^ and craving, the human study reported here is the first to explore the nature of the relationship within a dyad of peers, and to correlate it with cocaine consumption. It is also the first to find a direct effect of peer familiarity on drug consumption levels.

Some study limitations need to be acknowledged. First, the design of the rat study was experimental while the human study was observational and cross-sectional and collected retrospective information on episodes of stimulant consumption. In the latter, the outcome may be subject to recall and social desirability biases and to confounding. However, the approach we employed to question participants about their most recent episodes of drug intake is widely used to minimize recall bias. It is also used in many scales exploring behaviors, for example the Opiate Treatment Index, whose validity has been demonstrated in recent and less recent research^44^. Second, the use of a monthly measure of cocaine use would have prevented us from being able to study the presence of a peer, as cocaine use may have varied across the different episodes over a month. For this reason, to us, measuring use only for the most recent episodes was the only reliable way to simultaneously record the frequency of cocaine use and the relationship with a peer, if present.

It is possible that the lower relative risk of cocaine use frequency found in humans in presence of a peer could be attributable to drug sharing. While only data on syringe sharing was available and was not associated with the outcome, it is worth noting that it is unlikely that people decide to use less drugs in order to share what they have with an unfamiliar peer. Furthermore, we analyzed data using Poisson regression models (mixed model for rats, GEE for humans). In the human study we adjusted for possible confounders, in particular those known to be associated with frequency of cocaine use, such as unstable housing. Despite these limitations, we found very similar estimates for the association between frequency of consumption and the presence or the familiarity with a peer in both models, as the graphs clearly show. We realize that it is difficult to compare results in two different settings (experimental on animals versus observational in humans), however we took into account all social and behavioral variables, which can influence cocaine intake in humans. Only variables significantly associated with the outcome were considered and introduced into the final models for humans.

In conclusion, this study’s results, by showing parallel influences of proximal social factors on the frequency of drug consumption in rats and humans, highlight translation potential from rats to humans. The need for translational studies ^45,46^ is essential for a better understanding of the role played by social factors in addiction ^47^ and forces us to search for new models and solutions to translate as many aspects of behavior as we can.

Peer presence, peer familiarity and history of drug use, all have major effects on drug consumption. To better understand social influence mechanisms in drug addiction, research must now examine the neurobiological substrate of these observations.

Understanding how proximal social factors modulate drug consumption will help in the design of novel preventive and therapeutic strategies including social interventions to target drug-using populations. Furthermore, the presence of a non-familiar and possibly drug-naive peer would appear to be a driver for diminished stimulant intake. Given that there is still space for improvement in the management of cocaine-related disorders, these results may be crucial to develop harm reduction strategies for stimulant users. At the clinical level, this would translate into involving peers in treatment education. At the health policy level, it would mean promoting the use of harm reduction strategies, such as peer education on injection and the deployment of supervised consumption rooms.

## METHODS

### Human study

#### Design

The human study (DDYADS) is a cross-sectional survey implemented between October 2015 and June 2016 in 5 cities in France characterized by high prevalence of illicit drug use (Marseille, Paris, Montreuil, Saint Denis, Nice).

#### Participants

Seventy-seven French-speaking regular stimulant users – defined as using cocaine or methylphenidate ≥5 times a month – were recruited in different sites, including methadone centers, harm reduction centers, low-threshold mobile health units, and through word-of-mouth referrals, between October 2015 and June 2016. Non-prescribed methylphenidate was also considered as cocaine users may switch from cocaine to methylphenidate and vice-versa in the areas where the study was conducted, depending on black market availability and costs. The study received authorization from the national French Data Protection Authority (CNIL) and Aix-Marseille University’s institutional review board. All participants provided written informed consent.

#### Data collection

Data were anonymously collected through a face-to-face standardized questionnaire administered by trained interviewers. Interviews were conducted in a dedicated room at centers or in a café and lasted from 30 to 60 minutes. Participants were remunerated with a €15 gift voucher for completing the interview.

To minimize recall bias, participants retrospectively described episodes during the previous month where they used stimulants.

Social environment at the moment of stimulant use was described as follows: alone, with one peer, with a group (i.e., 2 or more peers). For episodes involving the participant and one peer, information on the peer was collected.

Peers were considered “familiar” if they were close friends or relatives, and if the participant could speak about his/her intimate life with them. Otherwise, peers were considered “non-familiar”. Participants were considered subordinate if they were economically dependent on the peer or if the peer was the leader in terms of drug use contexts (e.g. paying for the drug). Each drug use episode was characterized as follows: principal route of administration (intravenous, intranasal), type of stimulant (cocaine or methylphenidate), drug effect perception (from 1 to 5), concomitant use of other psychoactive substances including alcohol, location where episode took place (public versus private), state of mind (positive versus neutral versus negative), number of times drugs were consumed (including alcohol), and episode duration. We also collected data on participant characteristics including age, gender, employment status, educational level, housing situation (stable versus unstable), hazardous alcohol use (AUDIT-C score) (Bush & al., 1998), financial problems, including economic dependence on the peer, and the number of days the participant had used stimulants in the previous month.

#### Outcome

The outcome was the number of times drugs were consumed in one hour during each episode (i.e. frequency of drug intake).

The frequency of use during one episode (standardized by the duration of the episode) is an interesting measure, especially among cocaine users where the half-life of the drug may require repeated intake. From a public health viewpoint, this frequency is associated with several health risks (e.g. overdoses and other fatal events, as well as unsafe sexual behaviors^48–50^.

#### Statistical analysis

We considered each drug intake episode reported during the interview as a statistical unit. In order to take into account the within-subject correlation due to repeated measures (i.e., drug use episodes in the previous month) reported by the same individual, we used the Poisson Generalized Estimated Equation (GEE) approach for count data (Liam et al, 1986). First, we conducted a invariable analysis to test each variable describing the “social context”- peer presence (alone vs. one peer vs. group), familiarity, dominance of the peer – and potential correlates/confounders.

Second, based on the results of the univariable analysis, two models were built. In the first, we tested the role of peer presence (alone, with one peer, with a group i.e., two or more peers) on the frequency of stimulant use, after taking into account potential correlates/confounders including: 1) context of the stimulant use episode: type of location (private versus public place), route of administration (intravenous, intranasal), state of mind (positive, neutral, negative), number of other substances concomitantly used (including alcohol); 2) participant characteristics: gender, age, educational level (< high school certificate versus ≥ high school certificate), employment status, stable housing (i.e. renter or owner of their personal housing versus other), financial difficulties, hazardous alcohol use (AUDIT-C ≥4 for men and ≥3 for women), and number of days of stimulant use during the previous month (daily stimulant user versus other).

The second model was built to examine the role of peer familiarity (alone, with one familiar peer, with one non-familiar peer, with a group) on the frequency of drug intake after taking into account the potential correlates/confounders described in the first model.

We used a liberal p-value<0.20 in the univariable analysis to identify social context explanatory variables eligible for each multivariable model. A backward selection procedure was used to determine the two final multivariable models. We set the p-value threshold at α= 0.05 for these latter. All incidence rate ratio (IRR) estimates were reported with their 95% confidence intervals (95%CI) and tests were two-sided. STATA/SE version 12.1 software for Windows was used for the analyses.

### Rat Study

#### Animals & surgery

In the present study, 28 male Lister Hooded rats (Charles River Laboratories, Saint-Germain-sur-l’Arbresle, France) were housed in pairs upon their arrival. Rats were handled 2-3 times a week. They were maintained under 12-h light/dark cycles and had ad libitum access to food and water. All animal care and use conformed to the French regulation (Decree 2010-118) and were approved by local ethic committee and the University of Aix-Marseille (#3129.01). Using standard surgical procedures, silicon catheters were inserted into the right jugular vein of the rats. They exited dorsally between the scapulae. Further information on surgery and the apparatus used are provided in Supplement 1.

#### Apparatus

The experiment was conducted in four custom-built self-administration (SA) chambers (60 cmx 30 cmx 35 cm) made of opaque Perspex and divided into two compartments, separated by a grid. For the first experiment, only one of the two compartments was equipped with 2 chains and a stimulus light located on the right-hand wall. For the second experiment, both compartments were equipped so that both rats could self-administer. The grid allowed each rat visual, auditory and olfactory communication, limited tactile contact with its peer, and prevented each rat from accessing the tethering system of its peer. Drug infusions were delivered via intrajugular route of administration through tubing protected by a stainless steel spring, connected to 10 ml cocaine syringes positioned on motorized pumps (Razel Scientific Instruments, St. Albans, V T, USA) outside of the chamber.

All the chambers and pumps described above were controlled by a custom-built interface and associated software (built and written by Y. Pelloux).

#### Experimental Procedure

Two groups of rats (N=14 in each) were individually trained to pull a chain to self-administer cocaine (80µg per 90µl infusion in 5s) under a continuous schedule of reinforcement (Fixed Ratio 1 (FR1), 1 chain pulling results in 1 cocaine injection) for daily one-hour sessions. Cocaine was randomly assigned to one or the other of the two chains (the active chain). Pulling the active chain switched on the cue-light, delivered the cocaine to the blood stream and started a 20-s “time-out” during which any further pulling was recorded as perseveration, but had no other consequence. Pulling the other chain (inactive chain) was also recorded, as an error, and had no consequence. Once consumption became stable, the last 5 days of acquisition were used as a baseline for cocaine consumption when the rats were alone in the apparatus (condition “alone”, N=14 in each experiment). The rats were then exposed to 4 different self-administration conditions for 5 consecutive days each:

1) Peer Presence: In the presence of another rat (hereafter “peer”) having no access to cocaine. This peer could be either familiar (i.e. a cage-mate also trained for self-administration; N=8 in experiment 1 and N=6 in experiment 2) or a stranger (hereafter “non-familiar peer”) (i.e. a rat trained for cocaine self-administration but living in a different home-cage; N=6 in experiment 1 and N=8 for experiment 2). Peers were introduced into the cage after they had a minimum of 4 hours of abstinence from cocaine. In experiment 2, the peer introduced also had access to the drug.

The same familiar and non-familiar peers were used for each rat for all behavioral sessions.

2) Post-peer presence: Rats were “back alone” after exposure to peers (N=14 in each experiment) in order to assess whether peer presence could influence cocaine intake in future “alone” sessions.

3) In experiment 1 only: Non-familiar and cocaine-naive peer presence: rats from another group that had never been exposed to cocaine (N=11).

4) In experiment 1, subordinate/dominant peer presence: for the social interaction and CPP studies, dominant status within each pair of rats was assessed by behavioral observation during the first conditioning session of the CPP experiment. The number of pins and pounces for each rat was recorded during a 15-minute period of interaction. The “dominant” rat was assumed to be the one doing the most pinning and pouncing. The other rat was qualified as “subordinate”.

#### Outcome

For each one-hour behavioral session, the number of cocaine injections was recorded.

#### Statistical analysis

We assessed the association between the number of cocaine injections during the one-hour cocaine self-administration session (outcome; i.e. frequency) and the nature of the social relationship (i.e., familiar, not familiar) with rat peers, the history of cocaine exposure of rat peers (naive vs non-naive) and whether the rat peer was dominant or subordinate. Poisson mixed models were used to take into account the correlation over time between repeated measures of the outcome.

The following four models were analyzed, each including a random effect on time (in days), and the following experimental factor:

A) Peer presence: alone or with a non-naive peer, irrespective of the relationship (i.e. familiar/not familiar)

B) Familiarity: alone, with familiar peer, with non-familiar peer (only non-naive peers)

C) History of cocaine exposure: alone, with naive peer, with non-naive peer, back alone

D) Social status: dominant or subordinate (only non-naive familiar peers)

Statistical significance was set at α=0.05. All incidence rate ratio (IRR) estimates were reported with their 95% confidence intervals (95%CI) and tests were two-sided. STATA/SE version 12.1 software for Windows was used for the analyses.

#### Data availability statement

The datasets generated during and analyzed during the current study are available from the corresponding author on reasonable request.

## Acknowledgements

Our thanks to Prof. Jean-Paul Moatti for stimulating discussion at the start of the project, to Dany Palleressompoulle and Joel Baurberg for their technical support, as well as to the animal facilities’ personnel. Our thanks also to Camelia Protopopescu and Chiara Calzolaio for help with statistical analyses and to Jude Sweeney for the English revision and editing of the manuscript.

